# Development of photoactivatable drugs enables nicotinic optopharmacology

**DOI:** 10.1101/260232

**Authors:** Sambashiva Banala, Matthew C. Arvin, Nicolas M. Bannon, Xiao-Tao Jin, Yong Wang, Guiqing Zhao, John J. Marshall, Kyle R. Gee, David L. Wokosin, Anis Contractor, Henry A. Lester, Yevgenia Kozorovitskiy, Ryan M. Drenan, Luke D. Lavis

## Abstract

Photoactivatable (‘caged’) pharmacological agents have revolutionized neuroscience but the palette of available ligands is limited. We describe a general method for caging tertiary amines using an unconventional quaternary ammonium linkage that is chemically stable and elicits a desirable red-shift in activation wavelength. A photoactivatable nicotine (PA-Nic) prepared using this strategy could be uncaged via 1- or 2-photon excitation, making it useful for optopharmacology experiments to study nicotinic acetylcholine receptors (nAChRs) in different experimental preparations and spatiotemporal scales.

---

Photoactivatable (‘caged’) compounds are essential for modern biology and remain the only tools available for modulation of native proteins with high spatiotemporal resolution. The development of new photoactivatable compounds that are chemically stable and activatable with or 2-photon illumination remains an important application of organic chemistry to the biosciences. The ability to uncage ligands for glutamate and γ-aminobutyric acid (GABA) receptors enabled seminal ‘optopharmacology’ studies on the biophysics and subcellular location of functional proteins within complex biological environments^1–3^. Caged ligands targeting other receptors are rare, however, which has prevented precise examination of additional protein classes. We developed a general strategy for preparing photoactivatable drugs through alkylation of tertiary nitrogen atoms to form photolabile quaternary linkages. The resulting photoactivatable nicotine (PA-Nic) exhibits ideal chemical and spectroscopic properties for interrogating endogenous nicotinic acetylcholine receptors (nAChRs) in brain tissue.

Many pharmacological agents cannot be caged using standard strategies because they lack obvious attachment sites *(e.g.*, CO2H, OH, NH) for photolabile groups. Four examples of ‘uncageable’ drugs include nAChR agonists nicotine (1) and PNU-282,987 (2), the opioid fentanyl (3), and the selective serotonin reuptake inhibitor escitalopram (**4, Fig. 1a**). A shared feature of these compounds is a tertiary nitrogen, a common motif in many pharmacological agents. We envisioned a general caging strategy involving covalent attachment of a coumarin^4–6^ photolabile group to form a quaternary ammonium salt. This approach has been employed by chemists to create photoactivatable tertiary amines including polymer initiators^7^, amino acids^8^, mustards^9^, and tamoxifen^10^, but has never been used in a biological context. We initially applied this strategy to nicotine (**1**), alkylating with coumarin **5** to yield photoactivatable nicotine (PA-Nic, **6**; **Fig. 1b**). PA-Nic releases nicotine upon UV illumination (365 nm) with a photochemical quantum yield (Φ_pc_) of 0.74% (**Fig. 1c, Supplementary Fig. 1a; Supplementary Note**), and shows excellent dark stability in aqueous solution (**Fig. 1d**). Formation of the quaternary center at the 4-position of the coumarin elicits an unexpected ~15 nm red-shift in absorption maxima of the coumarin cage in PA-Nic (λ_max_ = 404 nm, **Fig. 1e**), an excellent match for LED or laser light sources centered near 400 nm. We then applied this strategy to prepare quaternary ammonium photoactivatable derivatives of drugs **2-4**, which also gave clean photoconversion (**Supplementary Fig. 1b-d, Supplementary Note**) with similar Φ_pc_, dark stability, and red-shifted absorbance maxima (**Supplementary Fig. 1e-g**). We confirmed that the shift in λ_max_ was due to presence of the quaternary center (**Supplementary Fig. 1h,i**) and determined the major and minor byproducts as coumarins **12** and **13**, respectively (**Supplementary Fig. 1j**).

**Figure 1.**
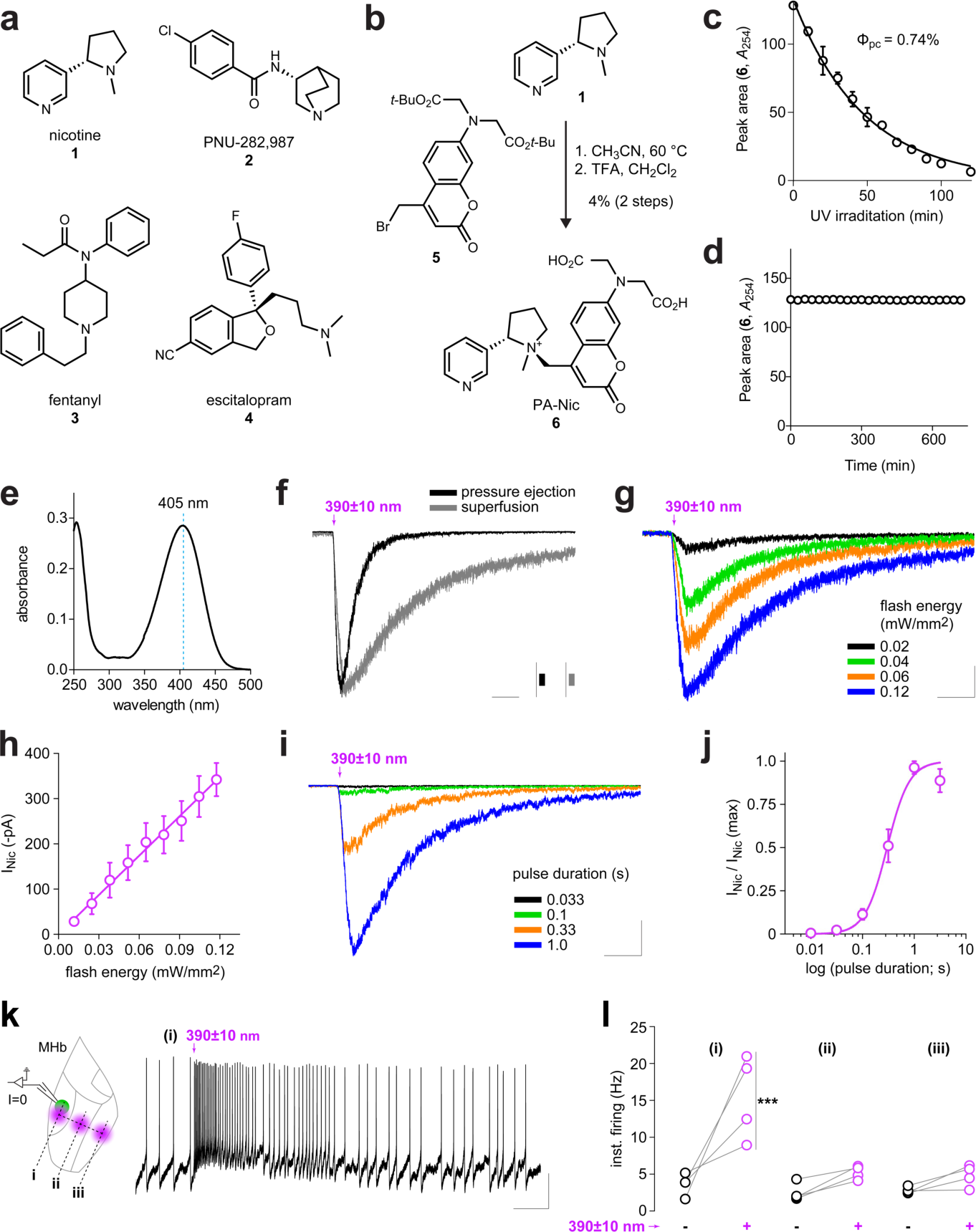
Development of PA-Nic and its utility for improved pharmacological studies of native nAChRs. (**a**) ‘Uncagable’ drugs **1**-**4**. (**b**) Synthesis of PA-Nic (**6**) (**c**) Plot of HPLC chromatogram peak area *vs.* irradiation time (365 nm) provides the photochemical quantum yield (ΦρΦ; solid line is exponential fit. (**d**) Dark stability of PA-Nic. (**e**) Absolute absorption spectrum of PANic (**6**, 20 μΜ). (**f**) Light-evoked currents following PA-Nic epi-illumination photolysis. Representative voltage clamp traces from individual MHb neurons are shown for different methods of PA-Nic application. Scale: 2.5 s, 100 pA (grey), 150 pA (black) (**g-j**) Controllable nicotine uncaging via PA-Nic epi-illumination photolysis. (**g**) Representative voltage clamp traces from a MHb neuron for light pulses (1 s) of varying intensity. Scale: 2.5 s, 75 pA (**h**) Resulting photochemical dose-response relation for peak currents (y=2.876x+2.1, R^2^=0.9921). Mean (±SEM, n=9 neurons) peak of light-activated currents plotted vs. input flash intensity. (**i**) Representative voltage clamp traces during light pulses of varying duration applied to a MHb neuron. Scale: 2.5 s, 250 pA (**j**) Graphical analysis of summary pulse duration data in **i**. The Hill equation was fitted to the data *(n_H_* (Hill slope)=2.0, duration at *^Λ^Α* max=0.3 s, R^2^=0.928) from n=5 neurons. (**k-l**) PA-Nic can be used to drive action potential firing via nAChRs. (**k**) MHb VI neurons were held in current clamp (I=0) configuration during epi-illumination photolysis of PA-Nic. A restricted field stop aperture permitted nicotine uncaging directly over the recorded VI neuron (i), or at 100 (ii) to 200 (iii) μm from the recorded cell. A representative trace is shown for a recording from a MHb VI neuron (right panel). Scale: 1.5 s, 15 mV (**l**) Before-after plots showing the peak action potential firing rate in individual MHb VI neurons (n=4 neurons from 2 mice) when nicotine was uncaged at position i, ii, or iii as indicated in **k**. *P* values (Bonferroni multiple comparison test following repeated measures two-way ANOVA): 0.0009 (i), 0.6627 (ii), >0.9999 (iii).

With the generality of this caging strategy established, we focused on PA-Nic (6), due to nicotine’s high selectivity for nAChRs and the societal and therapeutic significance of this molecule. We combined brain slice electrophysiology with epi-illumination flash photolysis to test the utility of PA-Nic and compare this compound to the known caged nicotine based on noncovalent complexes of ruthenium bis(bipyridine) (RuBi-Nic, **14, Supplementary Fig. 2a**).^11^ We prepared mouse brain slices containing the medial habenula (MHb), whose neurons express high levels of nAChRs^12^, co-release ACh and glutamate^13^, and regulate affective behavior and nicotine withdrawal^14^ (**Supplementary Fig. 2b-d**). Light flashes (1 s) evoked robust nicotinic currents when PA-Nic (80 μΜ) was superfused or applied locally via pressure ejection during voltage clamp recordings of MHb neurons (**Fig. 1f**). Photolysis of RuBi-Nic with a longer light dose (470 nm; 15 s) elicited a substantially smaller nicotinic current (**Supplementary Fig. 2e**). RuBi-Nic often appeared unstable when applied to brain slices via superfusion, resulting in uncontrollable/irreversible inward currents in the absence of light flashes (**Supplementary Fig. 2f**). In contrast to RuBi-Nic, PA-Nic photolysis does not occur with blue (470 nm) or green (560 nm) light flashes (100 ms, 0.06 mW/mm^2^; **Supplementary Fig. 2g**) enabling the use of other fluorophores or light-activated effectors.

Given the substantial improvement in performance over the existing RuBi-Nic, we further characterized PA-Nic in brain slice optopharmacology experiments. We confirmed that the main photochemical by-product of PA-Nic photolysis (**12**) was inert at nAChRs (**Supplementary Fig. 2h,i**). PA-Nic itself exhibited no antagonist properties at nAChRs, and our preparations of PANic were devoid of free nicotine (**Supplementary Fig. 2j,k**). The nAChR antagonist mecamylamine (mec) significantly reduced the inward current elicited by photolysis of PA-Nic (**Supplementary Fig. 2l**; pre-mec: 383 ± 102 pA; post-mec: 185 ± 51 pA; n=5; *p*=0.0359, paired i-test). Mecamylamine blocked ACh-evoked currents to a similar degree (pre-mec: 1027 ± 174 pA; post-mec: 464 ± 116 pA; n=5; *p*=0.0032, paired *t*-test) (**Supplementary Fig. 2m**). Finally, we measured the reversal potential of flash-activated currents by eliciting responses at holding potentials from -80 mV to 0 mV (**Supplementary Fig. 2n**). Inward currents activated by photolysis reversed at approximately +2 mV (**Supplementary Fig. 2o**), a characteristic feature of nAChRs.

Native nAChRs in brain slices are typically studied with bulk superfusion or pressure ejection of agonists. Unfortunately, bulk superfusion over-emphasizes the long-term effects of the drug (e.g., desensitization) and pressure ejection is limited by the difficulty associated with applying multiple drug concentrations to the same neuron while maintaining a stable recording. We tested whether photoactivation of PA-Nic would overcome these challenges. Epi-illumination light flashes elicited responses that increased in amplitude with increasing flash energy (**Fig. 1g,h**) and flash duration (**Fig. 1i,j**), with 1 s flashes typically eliciting the largest currents. Light-evoked release of nicotine allowed generation of complete photochemical dose-response relations in neurons, an advantage over application of single drug concentrations.

Caged pharmacological agents (e.g., glutamate, GABA) can be used to modulate action potential firing by precisely controlling ligand concentrations in brain slices with light. We tested whether PA-Nic superfusion and epi-illumination photolysis could be used during current clamp recordings to drive action potential firing via nAChR activation. Using a restricted field stop aperture (**Supplementary Fig. 2k**), we constrained the light flash to different ventral MHb subregions^15^ during current clamp recordings from neurons in the VI subregion (**Fig. 1k**). Brief flashes (33 ms) transiently enhanced firing when nicotine was uncaged over (within 0-60 μm, xy plane, location (i)) the recorded neuron, but elicited no change in firing when nicotine was uncaged over the VC (~100 μm, xy plane, location (ii)) or VL subregion (~200 μm, xy plane, location (iii)) (**Fig. 1k,l**). Two-way repeated-measures ANOVA of treatment (2; baseline vs. flash) x location (3; (i), (ii), (iii)) showed significant main effects of treatment [*F*(1,9)=21.07, *p*=0.0013], location [*F*(2,9)=19.79, *p*=0.0005], and a significant treatment x location interaction [*F*(2,9)=6.864, *p*=0.0155]. Bonferroni post-hoc testing confirmed a significant effect of photolysis only when nicotine was uncaged at location (i) (*p*=0.0009). PA-Nic, coupled with simple epi-illumination photolysis, could therefore be used for spatially-delimited modulation of action potential firing or presynaptic neurotransmitter release in diverse types of neurons that express nAChRs.

We tested the utility of PA-Nic under 2-photon (2P) activation to further probe the spatiotemporal precision of PA-Nic photolysis and determine whether the compound could be used in conjunction with 2-photon laser scanning microscopy (2PLSM). Slices were superfused with PA-Nic (100 μM) and photostimulation (720 nm) at multiple perisomatic locations evoked strong and repeatable inward currents (**Supplementary Fig. 3a**). Such currents could be evoked at wavelengths between 700 nm and 880 nm (**Supplementary Fig. 3b,c**), consistent with the properties of the coumarin caging moiety^4,6^. Mecamylamine superfusion significantly decreased the 2-photon uncaging response amplitudes (pre-mec: 11.06 ± 0.9 pA; post-mec: 8.56 ± 1.2 pA; n=11; *p*=0.008, paired *t*-test), and additional experiments using blockers of excitatory transmission ruled out any role for ionotropic glutamate receptors (data not shown). Importantly, the relatively narrow 2P-wavelength sensitivity of PA-Nic (<900 nm) circumvents cross-talk with many common fluorophores.

Next, we studied functional nAChRs in specific cellular compartments using 1-photon laser flash photolysis of PA-Nic. Using 2PLSM to visualize neuronal somata and dendrites, laser pulses (405 nm, ~1 μm spot diameter) were positioned adjacent to the soma during voltage clamp recordings. After confirming that moderate laser powers (1-2.5 mW, 50 ms) do not activate inward currents in the absence of PA-Nic (**Supplementary Fig. 3d,e**), we identified suitable stimulation parameters by first measuring the relationship between evoked inward currents versus laser power (**Supplementary Fig. 3f,g**) and versus pulse duration (**Supplementary Fig. 3h,i**). nAChR currents evoked by PA-Nic photolysis were antagonized by mecamylamine (**Fig. 2a**). nAChR activation could also be elicited from ventral tegmental area (VTA) neuronal somata using a local perfusion procedure (**Supplementary Fig. 3j,k**). VTA neurons exhibit modest nAChR expression compared to MHb neurons, demonstrating that PA-Nic can be used to study aspects of cholinergic transmission in multiple cell-types.

**Figure 2.**
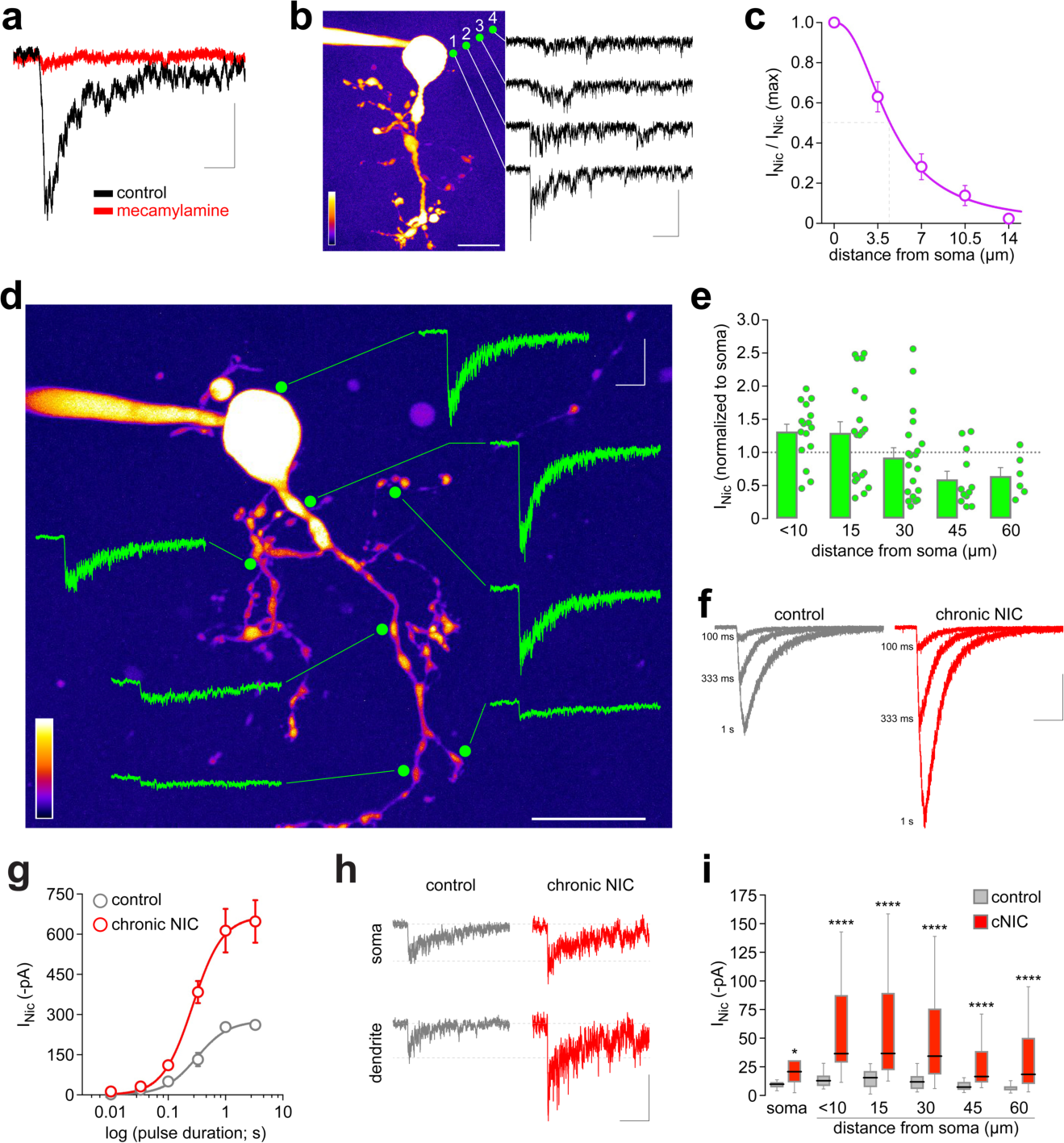
PA-Nic enables subcellular mapping of nAChRs following chronic nicotine treatment. (**a**) PA-Nic uncaging and mecamylamine antagonism. Nicotine was uncaged in a ~1 μm perisomatic spot with a 405 nm laser pulse (10 ms). Voltage-clamp currents are shown before (black trace) and 10 min after (red trace) mecamylamine (10 μM) superfusion. Scale: 50 ms, 100 pA (**b-c**) Lateral spread of uncaged nicotine, estimated electrophysiologically. (**b**) A representative 2PLSM image of a MHb neuron is shown. Nicotine was uncaged (green circles; 10 ms) at the surface (1; 0 μm), and at 3.5 (2), 7.0 (3), and 10.5 (4) μm from the cell surface. Representative traces from a single neuron are shown. Scales on image: (lower left = relative intensity; lower right = 20 μm). Scale (current): 500 ms, 30 pA (**c**) The Hill equation was fitted to the mean (±SEM) data (*n* (Hill slope)=2.293, R^2^=0.9074) from n=6 neurons, resulting in an estimate of 4.5 μm for the lateral distance at ½ max amplitude. (**d-e**) Subcellular mapping of nAChRs in MHb neurons. (**d**) Representative 2PLSM image of a cholinergic (ChAT(+)) MHb neuron, marked with uncaging positions (green circle) and the evoked response at that location. Scales: (lower left = relative intensity; lower right = 20 μm, upper right = 1 s, 60 pA) (**e**) Summary of position-dependent uncaging data for n=8 ChAT(+) MHb neurons using PA-Nic (80 μM). Nicotine uncaging responses were recorded at the soma and at dendritic locations at the indicated linear distance from the soma surface. The mean (+SEM) and individual responses are shown. (**f-i**) Interrogating chronic nicotine-mediated nAChR up-regulation with PA-Nic. (**f**) Representative traces are shown from a control and chronic nicotine-treated ChAT(+) MHb neuron stimulated via epi-illumination photolysis for the indicated durations. Scale: 200 pA, 2 s (**g**) The Hill equation was fitted to photochemical dose response data from control (7 neurons, 2 mice) or chronic nicotine-treated neurons (11 neurons, 3 mice) (control: *n*=1.7, duration at ½ max=0.3 s, R^2^=0.732; chronic nicotine: n=1.6, duration at *?* max=0.3 s, R^2^=0.89). (**h-i**) Chronic nicotine up-regulates nAChRs at all cellular locations tested. (**h**) Representative uncaging responses are shown from a control and chronic nicotine-treated neuron stimulated at the soma and at a dendrite ~30 μm from the soma using 405 nm laser photolysis (2 mW, 50 ms) of PA-Nic. Scale: 20 pA, 2 s (**i**) Box and whisker plots (whiskers=Tukey; black line=median; boxes=inner quartiles) of nicotine uncaging amplitudes at the indicated cellular location are shown for n=6 ChAT(+) neurons from n=3 control mice and n=14 ChAT(+) neurons from n=4 mice treated with chronic nicotine (cNIC). Mann-Whitney test: \**p*=0.0273, \*\*\*\**p*<0.0001.

Next, we estimated the spatial resolution of PA-Nic uncaging in MHb neurons. nAChR responses fell to 50% of their initial value at approximately 4.5 μm from the soma (**Fig. 2b,c**). We next determined whether functional nAChRs are present in dendrites of MHb neurons by uncaging nicotine at various dendritic locations. We found that nAChRs were functionally expressed in most subcellular locations, but the response amplitudes were greater near the soma and proximal dendrite compartments compared to distal dendrites (**Fig. 2d,e**). Nicotine appears in arterial blood rapidly (~10 s) in response to a puffed or vaped bolus^16^, and the close proximity (~20 μm) of neurons to capillaries implies that nicotine interacts with native nAChRs within seconds after reaching the brain. These results showcase the temporal and spatial precision afforded by laser flash photolysis of PA-Nic, which provides a pharmacokinetically relevant exposure paradigm superior to pressure ejection pulses.

Finally, we used PA-Nic epi-illumination and laser flash photolysis to study the pharmacology and cell biology of nAChR up-regulation, a key feature of human nicotine dependence. Chronic nicotine treatment results in up-regulation of nAChR function in MHb neurons^15^, including ChAT(+) neurons (**Supplementary Fig. 4a-d**), but it is unknown whether this up-regulation reflects a numerical increase in surface receptors or a shift in nAChR sensitivity. We used PA-Nic to generate a photochemical dose-response curve for nAChR currents in ChAT(+) MHb neurons from control and nicotine-treated mice, and found that chronic nicotine increased the pharmacological efficacy of acutely-applied nicotine without affecting its potency (**Fig. 2f,g**). This suggests the main action of chronic nicotine in MHb neurons may be to increase the overall number of functional surface receptors, without strongly affecting receptor sensitivity. Next, we used flash photolysis of PA-Nic combined with 2PLSM to examine how chronic nicotine exposure affects nAChR function in different cellular compartments. To do so, we examined laser flash-evoked nicotinic responses on somata as well as dendrites at various distances from the soma. Uncaging responses were enhanced in nicotine-exposed neurons, compared to control neurons, on both somata and dendrites (**Fig. 2h,i**). These data reveal that chronic nicotine exposure has a sensitizing effect on MHb neurons across the cell, potentially allowing ACh or nicotine to modulate key neuronal processes, such as action potential firing and dendritic integration. PA-Nic has thus afforded, to our knowledge, the first direct interrogation of postsynaptic nAChR plasticity mediated by chronic nicotine.

In summary, we have developed a strategy for preparing photoactivatable derivatives of previously uncagable drugs. The photoactivatable nicotine (PA-Nic, 6) developed using this approach efficiently releases nicotine (**Fig. 1b, Supplementary Fig. 1a**)—a specific nAChR agonist—making it advantageous over CNB-caged carbachol^17^, which activates all acetylcholine receptor types. PA-Nic can be activated by relatively short wavelength 1-photon (<470 nm) or 2-photon (<900 nm) light, allowing imaging in combination with other fluorophores. With the various modes of PA-Nic photolysis available, it is now possible to finely tune the spatiotemporal distribution of nicotine during optopharmacology experiments, allowing modeling of different aspects of nicotine exposure. PA-Nic could prove useful for studying cholinergic volume transmission *versus* point-to-point transmission ^18^. PA-Nic could be used in nAChR functional mapping^19^ and imaging experiments in key neuron types where dendritic and/or presynaptic Ca^2+^ dynamics are rapidly modulated by nAChR activation. Our initial mapping results (**Fig. 2d,e**) already suggest enhanced nAChR surface expression in proximal *versus* distal cellular compartments, and additional studies, such as those that account for dendritic filtering and differences in cellular surface area, will uncover further details of nAChR distribution in animals as a function of chronic nicotine. More generally, given the number of tertiary amine compounds in the pharmacopeia, the use of a photoscissile quaternary linkage should enable the development of other photoactivatable compounds to better model drug exposure and modulate native receptor proteins in the brain.

## Acknowledgments

We thank members of the Drenan and Lavis laboratories for helpful advice and discussion. This work was supported by the Howard Hughes Medical Institute (to S.B. and L.D.L.), National Institutes of Health (NIH) grants (DA035942 and DA040626 to R.M.D., MH099114 to A.C., DA037161 to H.A.L., NS054850 to D. James Surmeier), a PhRMA Foundation Fellowship (to M.C.A.), a Beckman Young Investigator Award (to Y.K.), a Bernice E. Bumpus Foundation Early Career Innovation Award (to. Y.K.), funding from the JPB Foundation, and funds from Northwestern University.

## Author Contributions

R.M.D., M.C.A., H.A.L., S.B., K.R.G., and L.D.L. conceived the project. M.C.A., N.M.B., D.L.W., X.J., Y.W., J.J.M., A.C., Y.K., R.M.D., S.B., and L.D.L. planned and/or executed experiments. D.L.W., Y.K. and K.R.G. contributed essential reagents and expertise. R.M.D., M.C.A., S.B., and L.D.L. wrote the paper with input from all authors. R.M.D. and L.D.L. supervised all aspects of the work.

## Competing Financial Interests

The authors declare no competing financial interests.

## Methods

**Chemical Synthesis and Photochemistry**. Experimental details, characterization for all novel compounds, and determination of photochemical byproducts of PA-Nic photolysis can be found in the Supplementary Note.

**UV-Vis Spectroscopy**. Spectroscopy was performed using 1-cm path length, 3.5-mL quartz cuvettes from Starna Cells or 1-cm path length, 1.0-mL quartz microcuvettes from Hellma. All measurements were taken at ambient temperature (22 ± 2 °C). Absorption spectra were recorded on a Cary Model 100 spectrometer (Agilent). Maximum absorption wavelengths (λmax) were taken in phosphate-buffered saline (PBS), pH 7.4.

**HPLC and LC-MS**. High performance liquid chromatography (HPLC) was performed on an Agilent 1200 Analytical HPLC system equipped with autosampler and diode array detector. To measure the photochemical quantum yield (**Fig. 1c**), the loss of PA-Nic (6) was monitored using a 4.6 × 150 mm Kinetex C18 column (Phenomenex) with a 5-95% gradient of CH_3_CN in H_2_O containing constant 0.1% v/v TFA. To examine the release of pharmacological agents **2-4** and coumarin byproducts from compounds **7-10 (Supplementary Fig. 1**), samples were run on tandem liquid chromatography-mass spectrometry (LC-MS) using an Agilent 1200 LC-MS system equipped with autosampler, diode array detector, and mass spectrometry detector using a 4.6 × 150 mm Gemini NX-C18 column with a 5-95% (**Supplementary Fig. 1a,d**) and 5-50% (**Supplementary Fig. 1b,c**) gradient of CH3CN in H2O containing constant 0.1% v/v TFA. **Photochemical Quantum Yield (Φ_pc_) Determination**. Photochemistry was performed in 1-cm path length/3.5 mL quartz cuvettes (Starna) in a Luzchem LZC 4V photoreactor equipped with 365 nm UV lamps, a carousel, and a timer as previously described^20^. Briefly, the light intensity was calibrated by potassium ferrioxalate actinometry. A solution of 60 mM K3Fe(C2O4)3 was irradiated using the photoreactor setup and released Fe^2+^ was determined by complexometry with 1,10 phenanthroline. Using the known photochemical quantum yield of this process (Φ_pc_ = 1.21), we determined the photon flux (I) = 3.57 × 10^−7^ ein/mincm^2^. For the conversion of PA-Nic (**6**) to nicotine (**1**), the samples were irradiated and a small aliquot (50 μL) was placed in an amber glass high recovery HPLC vial. These samples were analyzed by HPLC as described above. The photochemical quantum yield (Φ_PC_, mol/ein) was determined by fitting a plot of HPLC peak integral signal (S) vs. irradiation time to a one-phase exponential decay described by equation 1

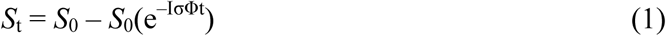

where *S*_0_ = signal prior to irradiation, t = irradiation time (min), *S*_*t*_ = signal at time t, I = irradiation (ein/mincm^2^), and σ = decadic extinction coefficient (in units of cm^2^/mol; 1000-fold higher than the ε value with units of M^−1^cm^−1^ based on cuvette geometry). For the conversion of compound **6** to compound **1**, we determined Φ_pc_ = 0.74% (**Fig. 1c**). Coumarin-based cages have found broad utility in release of small molecule modulators of biological activity ^4–6,21–27^We note the Φ_pc_ value for **6** is lower than coumarin caged molecules releasing better leaving groups such as carboxylates and phosphates^4,6,21,22^, but similar to the Φ_pc_ value of a 7-aminocoumarin caged compound releasing an amine via a carbamate linkage^6^. For other caged compounds, the photochemical quantum yield was determined by illumination with 405 nm LED (LOCTITE CL20 flood array) using 6 as a standard (**Supplementary Fig. 1e, Supplementary Note**). **Reagents for Neurobiology Experiments**. QX-314, CNQX, and D-AP5 were obtained from Tocris. RuBi-nicotine was obtained from Abcam. Acetylcholine (ACh), mecamylamine (mec), atropine, picrotoxin, and all other chemicals without a specified supplier were obtained from Sigma.

**Mice**. An animal study protocol pertaining to this study (#IS00003604) was reviewed and approved by the Northwestern University Institutional Animal Care and Use Committee. Procedures also followed the guidelines for the care and use of animals provided by the National Institutes of Health Office of Laboratory Animal Welfare. Mice were housed at 22 °C on a 12-hour light/dark cycle with food and water *ad libitum.* Mice were weaned on postnatal day 21 and housed with same-sex littermates. Unless stated otherwise, experiments were conducted on C57BL/6J mice obtained from Jackson Laboratories. ChAT-IRES-Cre^28^ (B6;129S6-*ChAT*^*tm2(cre)Lowl*^/J; Jax #006410) and Ai14^29^ (*B6.Cg-Gt(ROSA)26Sor*^*tm14(CAG-tdTomato)Hze*^*/*J; Jax #007914) mouse strains were crossed to yield ChAT-IRES-Cre::Ai14 mice, which express tdTomato in a Cre-dependent manner (i.e. in neurons with an active ChAT promoter). Expression of tdTomato in MHb ChAT(+) neurons was confirmed in these mice via immunohistochemistry and confocal microscopy (**Supplementary Fig. 2d**). All studies were restricted to male mice, age 8-24 wks.

**Chronic Nicotine Treatment**. Mice were treated with nicotine via drinking water as previously described^30^, with minor modifications. Nicotine hydrogen tartrate or L-tartaric acid (control group) were dissolved in tap water (pH 7.0) supplemented with saccharin sodium (3 mg/mL) to mask the bitter taste of nicotine. The following treatment schedule was used for nicotine (reported as nicotine free base) and tartaric acid, respectively (in μg/mL): Day 1-2 (50, 75), Day 3-4 (100, 150), Day 5 and beyond (200, 300). The latter doses were maintained by replacing drinking water solutions every 2-3 days, and mice were treated for at least 28 days prior to experimentation. We previously demonstrated up-regulation of nAChR function in MHb VI neurons using a different chronic nicotine exposure method: subcutaneous osmotic minipump implantation.^15^ Therefore, we validated this result with the nicotine drinking water method in C57BL/6 WT mice (**Supplementary Fig. 4a,b**) before conducting PA-Nic uncaging experiments coupled with chronic nicotine studies on ChAT-IRES-Cre::Ai14 mice (**Supplementary Fig. 4c,d** and **Fig. 2f-i**).

**Brain Slice Preparation**. For epi-illumination and 405 nm laser uncaging, brain slices were prepared as previously described^31^. Mice were anesthetized with Euthasol (sodium pentobarbital, 100 mg/kg; sodium phenytoin, 12.82 mg/kg) before trans-cardiac perfusion with oxygenated (95% O2/5% CO2), 4 °C N-methyl-D-glucamine (NMDG)-based recovery solution that contains (in mM): 93 NMDG, 2.5 KCl, 1.2 NH2PO4, 30 NaHCO_3_, 20 HEPES, 25 glucose, 5 sodium ascorbate, 2 thiourea, 3 sodium pyruvate, 10 MgSO_4_ 7H_2_O, and 0.5 CaCl_2_ 2H_2_O; 300-310 mOsm; pH 7.3-7.4). Brains were immediately dissected after the perfusion and held in oxygenated, 4 °C recovery solution for one minute before cutting a brain block containing the medial habenula and sectioning the brain with a vibratome (VT1200S; Leica). Coronal slices (250 μm) were sectioned through the MHb (**Supplementary Fig. 2b**) and transferred to oxygenated, 33 °C recovery solution for 12 min. Slices were then kept in holding solution (containing in mM: 92 NaCl, 2.5 KCl, 1.2 NaH2PO4, 30 NaHCO3, 20 HEPES, 25 glucose, 5 sodium ascorbate, 2 thiourea, 3 sodium pyruvate, 2 MgSO_4_ 7H_2_O, and 2 CaCl_2_ 2H_2_O; 300-310 mOsm; pH 7.3-7.4) for 60 min or more before recordings.

For 2-photon uncaging, brain slices were prepared as follows. Animals were deeply anesthetized by inhalation of isoflurane, decapitated, and the brain was rapidly removed and immersed in ice-cold oxygenated artificial cerebrospinal fluid (ACSF) containing (in mM) 127 NaCl, 2.5 KCl, 25 NaHCO3, 1.25 NaH2PO4, 2.0 CaCl_2_, 1.0 MgCl_2_, and 25 glucose (osmolarity ~310 mOsm/L). Tissue was blocked and transferred to a slicing chamber containing ice-cold ACSF, supported by a small block of 4% agar. Bilateral 250 μm-thick slices containing the MHb were cut on a Leica VT1000S and transferred into a holding chamber with ACSF equilibrated with 95% O2/5% CO2. Slices were incubated at 34 °C for 15-30 minutes prior to electrophysiological recording.

**UV-Vis Nicotine Uncaging**. Brain slices were transferred to a recording chamber being continuously superfused at a rate of 1.5-2.0 mL/min with oxygenated 32 °C recording solution. The recording solution contained (in mM): 124 NaCl, 2.5 KCl, 1.2 NaH_2_PO_4_, 24 NaHCO_3_, 12.5 glucose, 2 MgSO4-7H_2_O, and 2 CaCl_2_ 2H_2_O; 300-310 mOsm; pH 7.3-7.4). Picrotoxin (100 μM), CNQX (20 μM), and D-AP5 (50 μM) were added during subcellular uncaging experiments. TEA (5 mM) was added to the recording solution when the holding potential was 0 mV. Patch pipettes were pulled from borosilicate glass capillary tubes (1B150F-4; World Precision Instruments) using a programmable microelectrode puller (P-97; Sutter Instrument). Tip resistance ranged from 4.5 to 8.0 MΩ when filled with internal solution. The following internal solution was used (in mM): 135 potassium gluconate, 5 EGTA, 0.5 CaCl_2_, 2 MgCl_2_, 10 HEPES, 2 MgATP, and 0.1 GTP; pH adjusted to 7.25 with Tris base; osmolarity adjusted to 290 mOsm with sucrose. For subcellular uncaging, this internal solution also contained QX-314 (2 mM) for improved voltage control. We recorded from neurons in the ventral 50-60% of the MHb, as previously described^12,15^.

For epi-illumination uncaging experiments, a Nikon Eclipse FN-1 upright microscope equipped with infrared and visible differential interference contrast (DIC) optics and a 40X (0.80 NA) objective was used to visualize cells within brain slices. A computer running pCLAMP 10 software (Molecular Devices) was used to acquire whole-cell recordings along with an Axopatch 200B amplifier and a 16-bit Digidata 1440 A/D converter (both from Molecular Devices). Data were sampled at 10 kHz and low-pass filtered at 1 kHz. Immediately prior to gigaseal formation, the junction potential between the patch pipette and the superfusion medium was nulled. Series resistance was uncompensated. A light emitting diode (LED) light source (XCite 110LED; Excelitas) coupled to excitation filters (400/40 nm, 470/40 nm, and 560/40 nm bandpass) was used for photostimulation. Internal LEDs in the XCite 110LED were (center wavelength/FWHM, in nm): 385/30, 470/40, 560/80, 640/40. For near-UV photostimulation, flash wavelength was therefore approximately 390 ±10 nm. Light flashes were triggered by pCLAMP via TTL pulses. Flash energy output from the LED was determined by calibration using a photodiode power sensor (Model S120C; Thor Labs).

Some experiments involved recording inward currents following pressure ejection application of drug (ACh or PA-Nic) to the recorded cell using a drug-filled pipette, which was moved to within 20-40 μm of the recorded neuron using a micromanipulator. Using a Picospritzer (Parker Hannifin), a pressure ejection dispensed drug (dissolved in the same superfusion medium used on the slice) onto the recorded neuron. Ejection volume, duration, and ejection pressure varied depending on whether a short (ms) or long (s) pulse was required. The change in amplitude between the baseline and the peak was measured. Atropine (1 μM) was present in the superfusion medium when using ACh to prevent activation of muscarinic ACh receptors. To prevent stray light from prematurely uncaging nicotine inside the drug pipette, the field stop aperture on the microscope was restricted, and the drug pipette was retracted from the cell immediately following the application.

For focal nicotine uncaging using 1-photon laser (405 nm) flash photolysis, an Olympus BX51 upright microscope with a 60X (1.0 NA) objective was used to visualize cells. Prairie View 5 (Bruker Nano) software was used for acquisition via a Multiclamp 700B patch clamp amplifier (Molecular Devices). Analog signals were sampled at 5 kHz and low-pass filtered at 1 kHz, and an A/D converter (PCI-NI6052e; National Instruments) was used for digitization. Patch clamp recordings were carried out using the internal solution mentioned above, except that Alexa 568 (50 μM) or Alexa 488 (100 μM) was also included in the recording pipette to visualize cells using 2-photon laser scanning microscopy. After break-in, the internal solution with the Alexa dye was allowed to equilibrate for 15-20 min before imaging was initiated. The vast majority of 2PLSM 1-photon uncaging experiments utilized Alexa 488 and a Mai Tai HP1040 (Spectra Physics) tuned to 900 nm, whereas several pilot studies utilized Alexa 568 and a Mira 900 (Coherent) infrared laser (with Verdi 10W (532 nm) pump laser) tuned to 790 nm. The laser was pulsed at 80 MHz (Mai Tai HP) or 76 MHz (Mira) (<100 fs, ~250 fs pulse duration, respectively), and a Pockels cell (ConOptics) was used for power attenuation. The dual-channel, 2-photon fluorescence was detected by two non-de-scanned detectors; green and red channels (dual emission filters: 525/70 nm and 595/50 nm) were detected by the following Hamamatsu photomultiplier tubes (PMTs), respectively: end-on GaAsP (7422PA-40) and side-on multi-alkali (R3896). A 405 nm continuous wave laser (100 mW OBIS FP LX; Coherent) was used for photostimulation/uncaging via a second set of x-y galvanometers incorporated into the scanhead (Cambridge Technologies). 405 nm laser power was measured below the sample but above the condenser using a Field Master GS (LM10 HTD sensor head). A validation study on MHb neurons was conducted with the 405 nm laser in the absence of PA-Nic to identify any laser pulse-associated artifacts (current deflections) in the voltage clamp record. Small (<10 pA) inward currents were detected in several MHb cells when stimulated with 4-5 mW pulses (50 ms) (**Supplementary Fig. 3d,e**). No such currents were detected at <2.5 mW (**Supplementary Fig. 3d, e**), so laser pulses in this study did not exceed 2 mW.

PA-Nic was applied (40-80 μM in 5 mL) via superfusion to the slice using a recirculating perfusion system to conserve compound. Due to lower levels of nAChR expression compared to MHb, a higher concentration of PA-Nic (1 mM) was required to uncage appreciable nicotinic currents in VTA neurons. For such studies, local perfusion was performed; a glass pipette with a large diameter (30-40 μm) was back-filled with PA-Nic (1 mM) dissolved in ACSF, and was connected to a pressure ejection device. The PA-Nic-filled pipette was maneuvered to within ~50 μm of the recorded neuron and perfusion was triggered with a constant, low pressure (1-2 psi) (**Supplementary Fig. 3k, inset**). Prairie View 5 software was used to select spots in the field of view for focal uncaging of nicotine via 405 nm laser light flashes. Typical flash durations were 10-50 ms (unless otherwise stated), which were selected following an analysis of the relationship between pulse duration and nAChR activation (**Supplementary Fig. 3h,i**).

**2-Photon Nicotine Uncaging.** Slices were transferred to a recording chamber perfused with oxygenated ACSF at a flow rate of 2-3 mL/min. Whole-cell recordings were obtained from neurons of the MHb visualized under infrared Dodt contrast video microscopy using patch pipettes of ~2-5 MΩ resistance. One internal solution consisted of (in mM): 135 KMeSO_3_, 5 KCL, 0.5 CaCl2, 5 HEPES, 5 EGTA, 10 phosphocreatine disodium salt, 2 ATP, 0.5 GTP. A second internal solution, used in mecamylamine blockade experiments, contained (in mM): 120 CsMeSO4, 15 CsCl, 8 NaCl, 10 TEA-Cl, 10 HEPES, 2 QX-314, 4 ATP, 0.3 GTP, 0.2 EGTA. Alexa Fluor 488 (10-20 μM) was added to the internal solution to visualize cell morphology and confirm cell identity and location. Recordings were made using a Multiclamp 700B amplifier (Molecular Devices). Data were sampled at 10 kHz and filtered at 3 kHz, acquired in MATLAB (MathWorks). Series resistance, measured with a 5 mV hyperpolarizing pulse in voltage clamp, was under 30 MΩ and was left uncompensated. ACh sensitivity, observed in 100% of recorded neurons in the experiments validating PA-Nic uncaging efficacy, was tested using pressure ejection application of 1 mM ACh in ACSF.

Two-photon laser-scanning microscopy and two-photon laser photoactivation were accomplished on a modified Scientifica microscope with a 60× (1.0 NA) objective. Two mode-locked Ti:Sapphire lasers (Mai-Tai eHP Deep See and Mai-Tai eHP; Spectra Physics) were separately tuned, with beam power controlled by independent Pockels cells (ConOptics). The beams were separately shuttered, recombined using a polarization-sensitive beam-splitting cube (Thorlabs), and guided into the same galvanometer scanhead (Cambridge). The Mai Tai eHP Deep See was tuned to 910 nm for excitation of Alexa 488, and the Mai Tai eHP variably tuned between 690 and 1000 nm to uncage PA-Nic. A modified version of ScanImage was used for data acquisition^32^. PA-Nic was added by superfusion (100 μM), or via a Picospritzer. For PA-Nic pressure ejection, 300 ms pulses of 200 μM solution in ACSF were delivered at 5-10 psi through a patch pipette placed 20-60 μm away from the recorded cell. Successful photoactivation of PA-Nic was observed at the following parameter ranges: 5-20 ms pulse widths, 680-880 nm uncaging laser tuning, and 15-60 mW power measured at the sample plane. Two GaAsP photosensors (Hamamatsu, H7422) with 520/28 nm band pass filters (Semrock), mounted above and below the sample, were used for imaging Alexa 488 fluorescence signals.

**Immunohistochemistry and Confocal Microscopy**. Mice were anesthetized with sodium pentobarbital (100 mg/kg, i.p.) and transcardially perfused with 10 mL of heparin-containing phosphate buffered saline (PBS) followed by 30 mL of 4% paraformaldehyde. Brains were dissected, postfixed in 4% paraformaldehyde overnight at 4 °C, and dehydrated in 30% sucrose. Coronal brain slices (50 μm) were cut on a freezing/sliding microtome (SM2010R; Leica). MHb-containing slices were stained using the following procedure. Slices were first permeabilized for 2 min via incubation in PBST (0.3% Triton X-100 in PBS), followed by a 60 min incubation in blocking solution (0.1% Triton X-100, 5% horse serum in Tris-buffered saline (TBS)). Primary antibodies used in this study were as follows: goat anti-ChAT (Millipore; cat# AB144P), rabbit anti-DsRed (Clontech; cat# 632496). Primary antibodies were diluted in blocking solution (anti-ChAT at 1:100, anti-DsRed at 1:500). Slices were incubated in primary antibodies overnight at 4 °C. Three 5 min washes in TBST (0.1% Triton X-100 in TBS) were done before transferring slices to secondary antibodies for a 60-min incubation at room temperature. Secondary antibodies, diluted 1:500 in blocking solution, were as follows: anti-goat Alexa 488 (Invitrogen; cat# A11055), anti-rabbit Alexa 647 (Invitrogen; cat# A31573). Slices were washed as before, mounted on slides, and coverslipped with Vectashield. Staining in the MHb was imaged as previously described^12^ with a Nikon A1 laser-scanning confocal microscope.

**Statistics and Data Analysis**. Two-tailed statistical tests employed GraphPad Prism 6 software. For pharmacological experiments involving animal models, sample sizes were selected based on prior similar studies^12,15^, and mice of the appropriate age (8-24 wks) were randomly (informal randomization) chosen for assignment to specific experimental groups. For some studies involving recordings from neurons derived from mice treated with nicotine or control drinking water, the experimenter was blind to the treatment during data collection. Outlier data points were removed from data sets via the ROUT method (Q=1%) as previously described^33^.

**Data Availability**. The data that support the findings of this study, if not explicitly contained within the paper, Supplementary Information, or Supplementary Note, are available from the corresponding authors upon reasonable request.

---

**Supplementary Figure 1.**
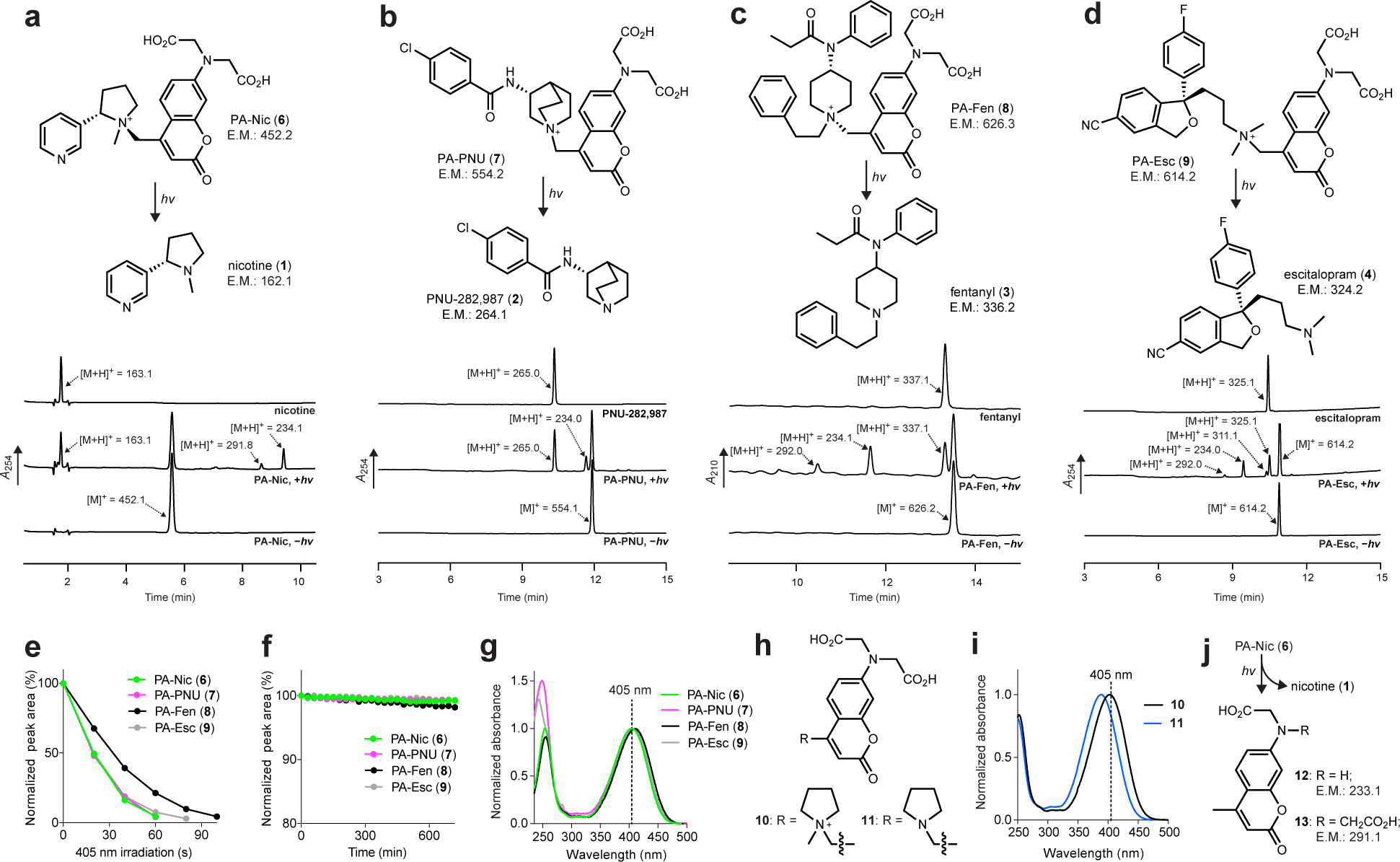
Synthesis and Characterization of Other Caged Pharmacological Agents. (**a-d**) Structures, exact mass (E.M.) and LC-MS traces of photoactivatable (PA) pharmacological agents and parent drug, before photolysis (-hv), and after photolysis (+hv); arrows show major ionic species from each peak. (**a**) PA-Nic (**6**) to nicotine (**1**). (**b**) PA-PNU (**7**) to PNU-282,987 (**2**). (**c**) PA-Fen (**8**) to fentanyl (**3**). (**d**) PA-Esc (**9**) to escitalopram (**4**). (**e**) Plot of normalized HPLC chromatogram peak area of PA compounds **6**-**9** *vs.* irradiation time (405 nm) to determine relative photochemical quantum yield(). (Φ_pc_) (**f**)Chemical (‘dark’) stability of PA compounds **6**-**9** (note scale) in the absence of light. (**g**) Normalized absorption spectrum of PA compounds **6**-**9**. (**h-i**) Quaternization of the coumarin cage elicits a 14 nm bathochromic shift in λ max. (**h**) Structures of model compounds **10** and **11**. (**i**) Normalized absorption spectra of **10** and **11**. (**j**) Chemical structure of coumarin photolysis byproducts **12** and **13** (**Supplementary Note**).

**Supplementary Figure 2.**
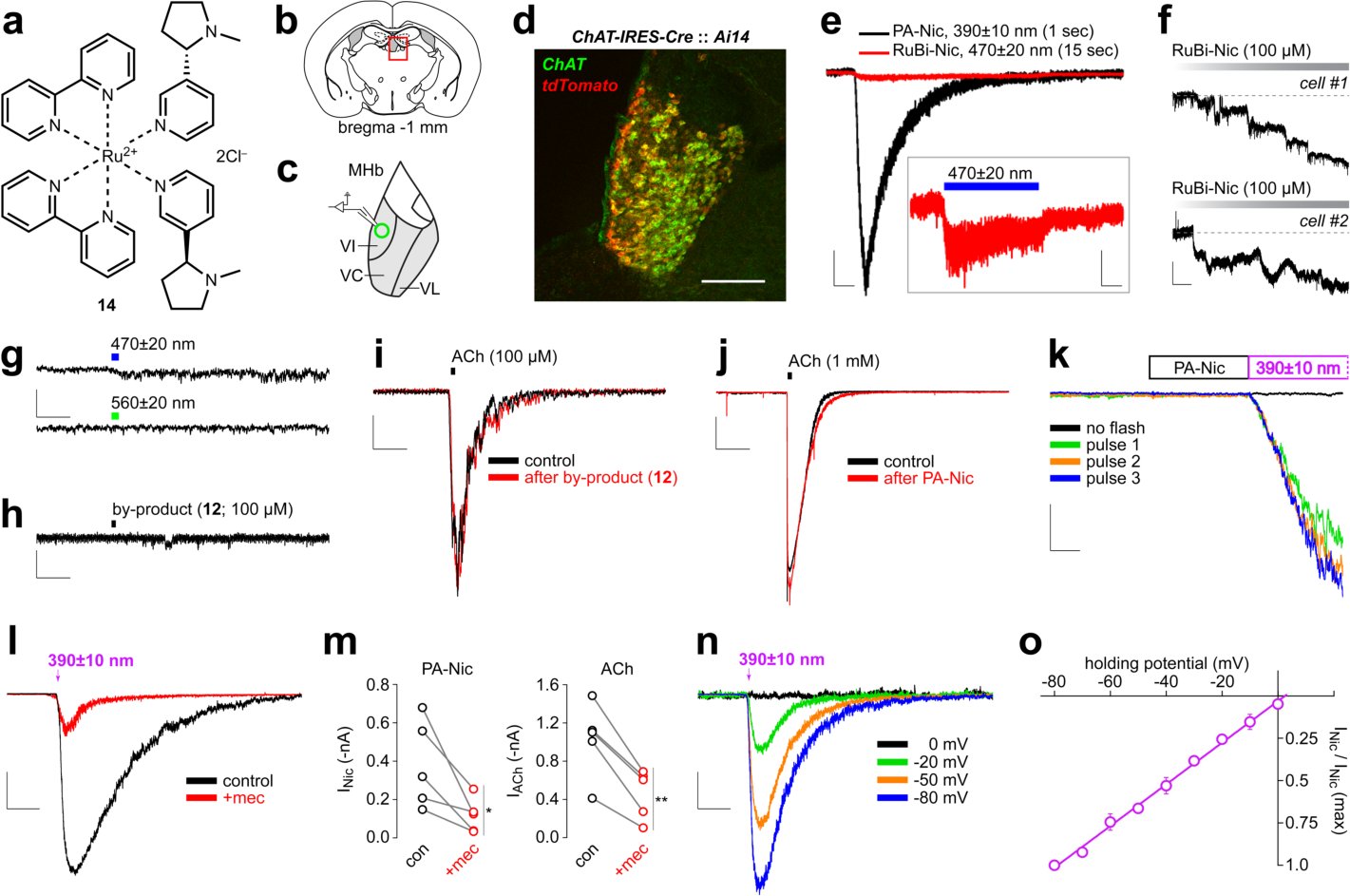
PA-Nic epi-illumination validation studies. (**a**) Chemical structure of RuBi-Nic (**14**). (**b**) Location of MHb in mouse brain near bregma -1 mm to -2 mm. (**c**) MHb subregions. Recordings were made from MHb neurons in the ventral inferior (VI) subregion. Other ventral MHb subregions: central (VC) and lateral (VL). (**d**) Validation of ChAT-IRES-Cre::Ai14 mice for targeted recordings from MHb cholinergic neurons. Coronal sections from ChAT-IRES-Cre::Ai14 mice containing MHb were costained with anti-ChAT and anti-DsRed antibodies. Scale: 175 μm. (**e**) Comparison of light-evoked responses using PA-Nic and RuBi-nicotine in MHb neurons. Epi-illumination flash PA-Nic (~405 nm, 1 s, 0.12 mW/mm^2^, 80 μM superfusion) and RuBi-nicotine (470 nm, 15 s, 0.12 mW/mm^2^, 1 mM local perfusion) uncaging responses are plotted on the same time and current scale. Scale bars: 2 s, 60 pA. Inset: RuBi-nicotine response re-plotted to show the small inward current deflection. Scale bars: 2 s, 5 pA (**f**) RuBi-nicotine is unstable in brain slices. Voltage clamp recordings before/during RuBi-nicotine (100 μM) superfusion for n=2 independent VTA cells are shown. No light flashes were delivered. Scale bars: 100 pA, 2 min. (**g**) No PA-Nic uncaging with blue or green light. PA-Nic was applied to a voltage-clamped MHb neuron, and blue (~470 nm) or green (~560 nm) light flashes (100 ms, 0.06 mW/mm^2^) were applied. Representative traces are shown for n=2 recordings. Scale: 500 ms, 5 pA. (**h-i**) The main photochemical by-product of PA-Nic photolysis does not act as an agonist or antagonist at nAChRs. (**h**) By-product (**12**), monoalkylcoumarin, was applied via pressure ejection to a voltage-clamped MHb neuron (100 μM, 12 psi, 125 ms). Scale: 1 s, 5 pa (**i**) ACh was applied to a voltage-clamped MHb neuron before (black trace) and after (red trace) superfusion of by-product (**12**; 100 μM). Scale: 3 s, 20 pA. (**j**) PA-Nic does not antagonize nAChRs in MHb neurons. ACh (1 mM) was applied to a voltage-clamped MHb neuron via pressure ejection before (black trace) and after (red trace) superfusion of PA-Nic (80 μM). Scale: 3 s, 250 pA (**k**) Repeated nicotine uncaging using PA-Nic pressure ejection. PA-Nic (100 μM) was applied locally to a MHb neuron via pressure ejection (500 ms, 12 psi), followed immediately by a light flash with the microscope field stop aperture fully restricted, for several trials. Scale bars: 150 ms, 100 pA (**l-m**) PA-Nic voltage clamp responses are antagonized by mecamylamine. (**l**) Representative voltage clamp traces are shown for light-evoked currents before (black trace) and 10 min after (red trace) superfusion of mecamylamine (10 μM). PA-Nic was applied locally to the cell via pressure ejection using a 1 s flash (0.12 mW/mm^2^). Scale: 1.5 s, 150 pA (**m**) Before-after plots for PANic light-evoked currents (left panel) are compared to ACh-evoked currents (right panel) in the same neuron. *P* values (paired t-test): 0.0359 (PA-Nic; n=5 neurons), 0.0032 (ACh; n=5 neurons). (**n-o**) PA-Nic light-evoked currents are voltage-dependent. (**n**) Representative light-evoked currents from the same neuron are shown for different holding potentials. PA-Nic was applied locally to the cell via pressure ejection using a 1 s flash (0.12 mW/mm^2^). Scale: 1.5 s, 200 pA (**o**) Current-voltage relation: currents during nicotine uncaging at various holding potentials. Data show mean ± extrapolates to a reversal potential of ~ +2 mV.

**Supplementary Figure 3.**
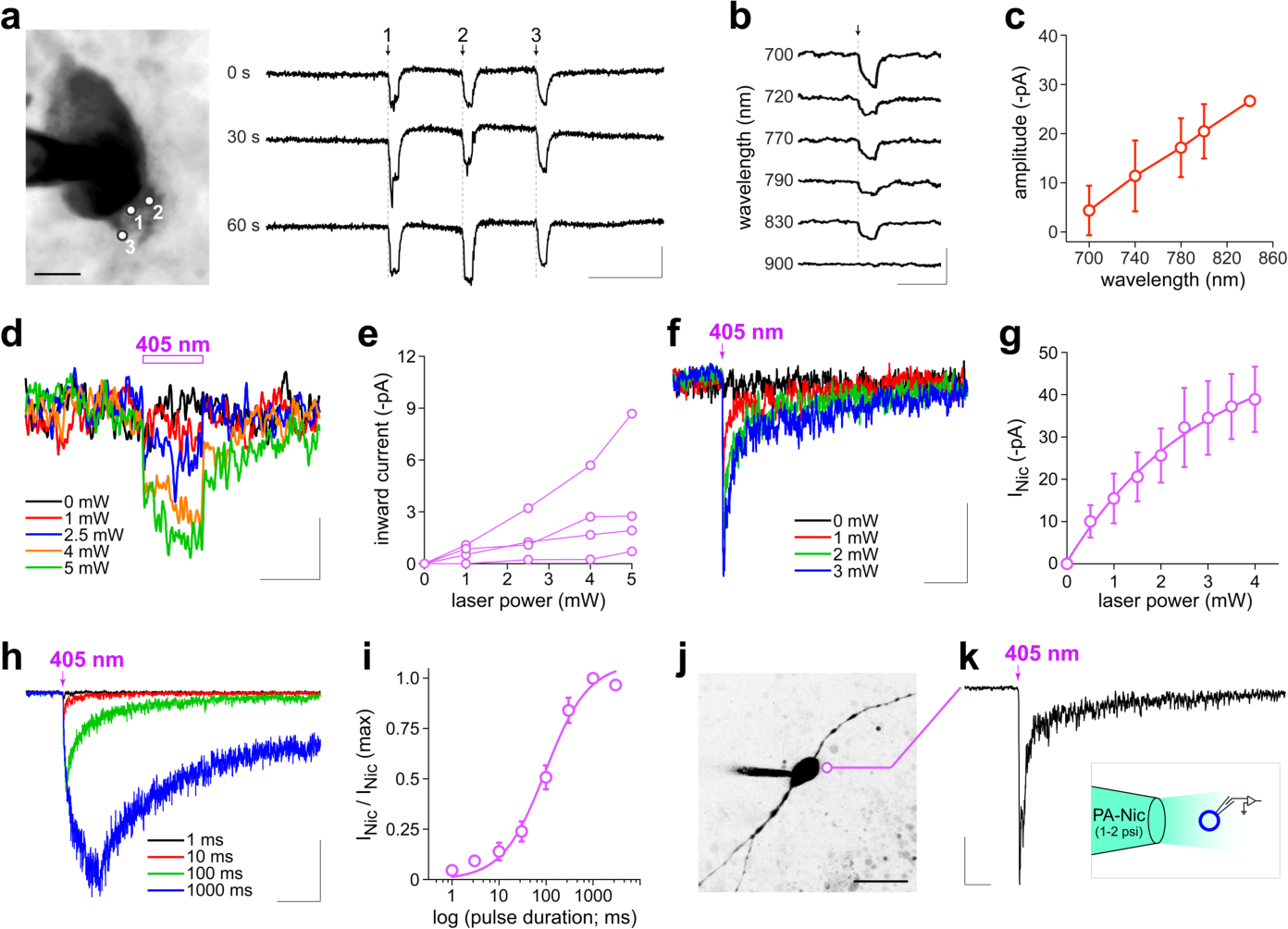
PA-Nic laser flash photolysis validation studies. (**a**) Representative 2-photon laser scanning microscopy image of a patch-clamped, dye-filled MHb neuron soma, including three perisomatic uncaging positions. Scale (image): 6 μm; Scale (current): 50 ms, 50 pA. Evoked currents (720 nm, 60 mW, 5 ms) are shown for each position (trials recorded at 30 s intervals). (**b-c**) 2-photon uncaging of nicotine is successful over a broad range of wavelengths. (**b**) At one location on a proximal dendrite, uncaging (60 mW, 10-20 ms) was successful at wavelengths between 700 and 830 nm, but failed to evoke responses at 900 nm. Scale: 50 ms, 50 pA (**c**) In one cell, five uncaging locations were tested for their wavelength dependence and current amplitude was correlated with increasing wavelength (Pearson r=0.6179, *p*=0.0107, R^2^=0.3818). (**d-e**) Small inward currents during pulses with high laser power. In the absence of PA-Nic, patch-clamped MHb neurons were stimulated with 405 nm laser flashes (50 ms, perisomatic uncaging position, n=4 neurons) using a range of laser powers. **d** shows average traces for n=4 neurons at the indicated laser power. Scale bars: 50 ms, 1.5 pA. **e** shows a plot of the peak inward current for these n=4 neurons at the indicated laser power. (**f-g**) Relationship between laser power and inward current for PA-Nic uncaging. PA-Nic (80 μM) was applied via superfusion to patch-clamped MHb neurons, and nAChR-mediated currents were evoked via 405 nm laser flashes (10 ms, perisomatic position, n=4 neurons) using a range of laser powers. **f** shows voltage recordings from a representative cell during PANic uncaging using the indicated laser power. Scale bars: 1 s, 16 pA. **g** shows average peak inward current for PA-Nic uncaging-evoked responses at the indicated laser power. A single-phase exponential function was fitted to the data (R^2^=0.537). (**h-i**) Relationship between laser pulse duration and inward current for PA-Nic uncaging. PA-Nic (80 μM) was applied via superfusion to patch-clamped MHb neurons, and nAChR-mediated currents were evoked via 405 nm laser flashes (1 mW laser power, perisomatic position, n=6 neurons) using a range of stimulation durations. **h** shows voltage recordings from a representative cell during PA-Nic uncaging using the indicated pulse duration. Scale bars: 1 s, 40 pA. **i** shows average (normalized to maximum for each cell) peak inward current at each pulse duration. The Hill equation was fitted to the data (n_H_ (Hill slope)=1.0, duration at ¼ max=100 ms, R^2^=0.943) from n=6 neurons. (**j-k**) PA-Nic uncaging in ventral tegmental area (VTA) neurons. (**j**) 2PLSM image of a patch-clamped, Alexa 488-filled VTA neuron is shown. Scale: 40 μm. (**k**) Laser flash-evoked nicotinic currents in VTA neurons. A representative (of n=3 neurons) inward current is shown in response to a 405 nm laser pulse (2 mW, 50 ms, perisomatic uncaging position). Scale: 250 ms, 100 pA. Inset: PA-Nic local perfusion was used for VTA uncaging.

**Supplementary Figure 4.**
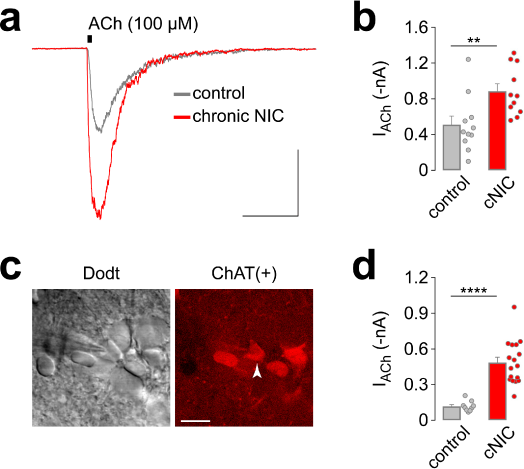
Up-regulation of nAChRs in MHb ChAT(+) neurons. (**a-b**) Chronic nicotine treatment up-regulates nAChRs in MHb neurons of C57BL/6 WT mice. (**a**) Representative ACh (100 μM)-evoked currents from a control or chronic nicotine-treated MHb neuron from C57BL/6 WT mice treated with control or chronic nicotine via nicotine-laced drinking water. Scale: 500 ms, 200 pA. (**b**) Bar/dot plot showing mean (+SEM) and individual responses to pressure-applied ACh (100 μM) in MHb neurons (# of neurons: n=11 control; n=11 nicotine) from control and chronic nicotine-treated mice (# of mice: n=2 control; n=2 nicotine). \*\**p*=0.0034 (Mann-Whitney test). (**c-d**) Chronic nicotine treatment up-regulates nAChRs in MHb ChAT(+) neurons. (**c**) ChAT-expressing MHb neurons in brain slices during patch clamp recordings. Representative Dodt contrast (left) and 2PLSM (right) image of tdTomato expression in ChAT(+) MHb neurons from ChAT-IRES-Cre::Ai14 mice (arrow) is shown. Scale: 20 μm. (**d**) nAChR functional up-regulation in ChAT(+) MHb neurons. ChAT-IRES-Cre::Ai14 mice (# of mice: n=2 control; n=3 nicotine) were treated with control or chronic nicotine for 4-6 weeks via their drinking water, and ACh (100 μM)-evoked currents were recorded from visually-identified ChAT(+) neurons (# of neurons: n=9 control; n=17 nicotine). \*\*\*\**p*<0.0001 (Mann-Whitney test)

## References

1. Matsuzaki, M., Ellis-Davies, G.C., Nemoto, T. et al., Nat. Neurosci. 4, 1086–1092 (2001).

2. Ellis-Davies, G.C., Nat. Methods 4, 619–628 (2007).

3. Matsuzaki, M., Hayama, T., Kasai, H. et al., Nat. Chem. Biol. 6, 255–257 (2010).

4. Furuta, T., Wang, S.S., Dantzker, J.L. et al., Proc. Natl. Acad. Sci. U.S.A. 96, 1193–1200 (1999).

5. Hagen, V., Dekowski, B., Kotzur, N. et al., Chem. Eur. J. 14, 1621–1627 (2008).

6. Hagen, V., Dekowski, B., Nache, V. et al., Angew. Chem. Int. Ed. 44, 7887–7891 (2005).

7. Sarker, A.M., Kaneko, Y., and Neckers, D.C., J. Photochem. Photobiol. A 117, 67–74 (1998).

8. Petersson, E.J., Choi, A., Dahan, D.S. et al., J. Am. Chem. Soc. 124, 12662–12663 (2002).

9. Mccarron, S.T., Feliciano, M., Johnson, J.N. et al., Bioorg. Med. Chem. Lett. 23, 2395–2398 (2013).

10. Asad, N., Deodato, D., Lan, X. et al., J. Am. Chem. Soc. 139, 12591–12600 (2017).

11. Filevich, O., Salierno, M., and Etchenique, R., J. Inorg. Biochem. 104, 1248–1251 (2010).

12. Shih, P.Y., Engle, S.E., Oh, G. et al., J. Neurosci. 34, 9789–9802 (2014).

13. Ren, J., Qin, C., Hu, F. et al., Neuron 69, 445–452 (2011).

14. Salas, R., Sturm, R., Boulter, J. et al., J. Neurosci. 29, 3014–3018 (2009).

15. Shih, P.Y., Mcintosh, J.M., and Drenan, R.M., Mol. Pharmacol. (2015).

16. Rose, J.E., Behm, F.M., Westman, E.C. et al., Drug Alcohol Depend. 56, 99–107 (1999).

17. Milburn, T., Matsubara, N., Billington, A.P. et al., Biochemistry 28, 49–55 (1989).

18. Parikh, V., Kozak, R., Martinez, V. et al., Neuron 56, 141–154 (2007).

19. Khiroug, L., Giniatullin, R., Klein, R.C. et al., J. Neurosci. 23, 9024–9031 (2003).

## Methods-Only References

20. Grimm, J.B., English, B.P., Choi, H. et al., Nat. Methods 13, 985–988 (2016).

21. Herbivo, C., Omran, Z., Revol, J. et al., Chembiochem 14, 2277–2283 (2013).

22. Shembekar, V.R., Chen, Y., Carpenter, B.K. et al., Biochemistry 46, 5479–5484 (2007).

23. Bourbon, P., Peng, Q., Ferraudi, G. et al., Bioorg. Med. Chem. Lett. 23, 2162–2165 (2013).

24. Gandioso, A., Cano, M., Massaguer, A. et al., J. Org. Chem. 81, 11556–11564 (2016).

25. Nadler, A., Yushchenko, D.A., Muller, R. et al., Nat. Commun. 6, 10056 (2015).

26. Schaal, J., Dekowski, B., Wiesner, B. et al., Chembiochem 13, 1458–1464 (2012).

27. Schoenleber, R.O. and Giese, B., Synlett, 501–504 (2003).

28. Rossi, J., Balthasar, N., Olson, D. et al., Cell Metab. 13, 195–204 (2011).

29. Madisen, L., Zwingman, T.A., Sunkin, S.M. et al., Nat. Neurosci. 13, 133–140 (2010).

30. Zhao-Shea, R., Degroot, S.R., Liu, L. et al., Nat. Commun. 6, 6770 (2015).

31. Engle, S.E., Broderick, H.J., and Drenan, R.M., J. Vis. Exp., e50034 (2012).

32. Pologruto, T.A., Sabatini, B.L., and Svoboda, K., Biomed. Eng. Online 2, 13 (2003).

33. Motulsky, H.J. and Brown, R.E., BMC Bioinformatics 7, 123 (2006).

